# Measuring experimental cyclohexane-water distribution coefficients for the SAMPL5 challenge

**DOI:** 10.1101/063081

**Authors:** Ariën S. Rustenburg, Justin Dancer, Baiwei Lin, Jianwen A. Feng, Daniel F. Ortwine, David L. Mobley, John D. Chodera

**Author notes:** Current address: Theravance Biopharma, South San Francisco, CA 94080, United States.

## Abstract

Small molecule distribution coefficients between immiscible nonaqueuous and aqueous phases—such as cyclohexane and water—measure the degree to which small molecules prefer one phase over another at a given pH. As distribution coefficients capture both thermodynamic effects (the free energy of transfer between phases) and chemical effects (protonation state and tautomer effects in aqueous solution), they provide an exacting test of the thermodynamic and chemical accuracy of physical models without the long correlation times inherent to the prediction of more complex properties of relevance to drug discovery, such as protein-ligand binding affinities. For the SAMPL5 challenge, we carried out a blind prediction exercise in which participants were tasked with the prediction of distribution coefficients to assess its potential as a new route for the evaluation and systematic improvement of predictive physical models. These measurements are typically performed for octanol-water, but we opted to utilize cyclohexane for the nonpolar phase. Cyclohexane was suggested to avoid issues with the high water content and persistent heterogeneous structure of water-saturated octanol phases, since it has greatly reduced water content and a homogeneous liquid structure. Using a modified shake-flask LC-MS/MS protocol, we collected cyclohexane/water distribution coefficients for a set of 53 druglike compounds at pH 7.4. These measurements were used as the basis for the SAMPL5 Distribution Coefficient Challenge, where 18 research groups predicted these measurements before the experimental values reported here were released. In this work, we describe the experimental protocol we utilized for measurement of cyclohexane-water distribution coefficients, report the measured data, propose a new bootstrap-based data analysis procedure to incorporate multiple sources of experimental error, and provide insights to help guide future iterations of this valuable exercise in predictive modeling.

**Abbreviations used in this paper:**

SAMPL: Statistical Assessment of the Modeling of Proteins and Ligands
log P: log _10_ partition coefficient
log D: log _10_ distribution coefficient
LC-MS/MS: Liquid chromatography - tandem mass spectrometry
HPLC: High-pressure liquid chromatography
MRM: Multiple reaction monitoring
DMSO: Dimethyl sulfoxide
PBS: Phosphate buffered saline
RPM: Revolutions per minute
CV: Coefficient of variation
MAP: Maximum *a posteriori*
MCMC: Markov chain Monte Carlo

## I. INTRODUCTION

Rigorous assessment of the predictive performance of physical models is critical in evaluating the current state of physical modeling for drug discovery, assessing the potential impact of current models in active drug discovery projects, and identifying limits of the domain of applicability that require new models or improved algorithms. Past iterations of the SAMPL (**S**tatistical **A**ssessment of the **M**odeling of **P**roteins and **L**igands) experiment have demonstrated that blind predictive challenges can expose weaknesses in computational methods for predicting protein-ligand binding affinities and poses, hydration free energies, and host-guest binding affinities [1–4]. In addition, these blind challenges have contributed new, high-quality datasets to the community that have enabled retrospective validation studies and data-based parameterization efforts to further advance the current state of physical modeling.

By focusing community effort on the prediction of hydration free energies in the first few iterations of this challenge, the SAMPL experiments have now brought physical modeling approaches to the point where they can reliably identify erroneous experimental data [5]. While hydration free energy exercises have shown their utility in improving the state of physical modeling, they are laborious, require specialized equipment no longer found in modern laboratories, are (at least using traditional protocols) limited in dynamic range, and are of questionable applicability in their ability to mimic protein-to-solvent transfer. As a result, no experimental laboratory has emerged to provide new hydration free energy measurements to sustain this aspect of the SAMPL challenge. We sought to replace this component of the SAMPL challenge portfolio with a new physical property that was easy to measure, accessible to multiple laboratories, had a wide dynamic range (in a free energy scale), and better mimicked physical and chemical effects relevant to protein-to-solvent transfer free energies, but was still free of the conformational sampling challenges protein-ligand binding affinities present. As the measurement of partition and distribution coefficients is now widespread in pharma (due to its relevance in optimizing lipophilicity of small molecules), we posited that a blind challenge centered around the prediction of distribution coefficients—which face many of the same physical and chemical effects (such as protonation state [6, 7] and tautomer issues [8]) observed in protein-ligand binding—might provide such a challenge.

While the measurement of octanol/water distribution coefficients is commonplace (a 2008 benchmark of structure-and property-based log P prediction methods used 96,000 experimental measurements [9]), a number of previously-reported complications in the physical simulation of 1-octanol suggested that this might be too complex for an initial distribution coefficient challenge [10–13], despite some recent reports of success [14]. In particular, water-saturated octanol is very wet, containing 47±1 mg water/g solution [15], and forms complex microclusters or inverse-micelles that create a heterogeneous environment that persist for long simulation times [10–13]. For the inaugural distribution coefficient challenge in SAMPL5, we therefore chose to measure cyclohexane/water distribution coefficients. The water content of water-saturated cyclohexane is much lower than water-saturated octanol—0.12 mg water/g solution, approximately 400 times smaller [16–18], and possesses no long-lived heterogeneous structure [19].

The number of freely available sources of cyclohexane-water partition is very limited, and for the purpose of the SAMPL5 distribution coefficient challenge[20], blind data was required. As part of an internship program at Genentech arranged by the coauthors, the lead author was dispatched to work out modifications of a high-throughput shake-flask protocol [21] currently in use for octanol/water distribution coefficient measurements. In particular, the low dielectric constant of cyclohexane (2.0243) compared to 1-octanol (10.30) [22] and cyclohexane’s surprising ability to dissolve laboratory consumables presented some unexpected challenges. In this report, we describe the modified protocol that resulted, and provide suggestions on how it can further be refined for future iterations of the distribution coefficient challenge. Of 95 lead-like molecules with diverse functional groups selected for measurement, we report 53 log D measurements that passed quality controls that were used in the SAMPL5 challenge.

To ensure the reported experimental dataset is useful in assessing, falsifying, and improving computational physical models of physical properties, we require a robust approach to estimating the experimental error (uncertainty in experimental measurements). We explored several procedures for propagating known sources of error in the measurement process into the final reported log distribution coefficients, and report those efforts here. Our primary approach features a parametric bootstrap, which allows the use of a physical model of the data generating process to sample additional realizations of the data, using distributions specified in the model. These additional realizations are new data points, over which estimates can be calculated. We compared this to a nonparametric bootstrap, which can be useful if a physical model can not be constructed. This method generates new data points as well, but it constructs them from selection with replacement from the existing data. We also calculated the arithmetic mean and standard error of the measured data. We hope that future efforts to measure cyclohexane-water distribution coefficients can benefit from the model we have developed, so that this work will also be useful for future challenges.

All code used in the analysis, as well as raw and processed data, can be found at https://github.com/choderalab/sampl5-experimentallogd-data

### Theory of distribution coefficients

The *distribution coefficient,* D, is a measure of preferential distribution of a given compound (solute) between two immiscible solvents at a specified pH, usually specified as log D in its base-10 logarithmic form,

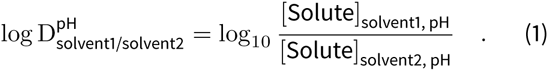

Typically, one solvent is aqueous and buffered at the specified pH (e.g. Tris pH 7.4), while the other is apolar (e.g. 1-octanol). At the given pH, the solute may populate multiple protonation or tautomeric states, but the total concentration summed over all states is used in the calculation of concentrations in Equation (1). The total salt concentration of the aqueous phase can also play a role, in case salts can provide stabilization of an ionic state of the ligand in the aqueous phase [23]. Additionally, temperature can cause shifts in the equilibrium populations [23]. Because of this, care must be exercised when comparing distribution coefficients obtained under different experimental conditions.

For the SAMPL5 challenge, we concern ourselves with the cyclohexane-water distribution coefficient, where phosphate-buffered saline (PBS) at pH 7.4 is used for the aqueous phase, at a temperature of 25°C:

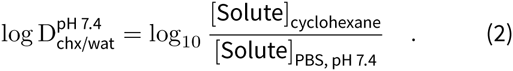

Another commonly reported value is the *partition coefficient P*, which quantifies the relative concentration of the neutral species in each phase, again usually specified in log_10_ form,

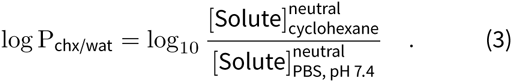

For ligands with a single titratable site and known p*K*_*a*_, one is can readily convert between log P and log D for a given pH iso (see, e.g. [23]), but ligands with more complex protonation state effects or tautomeric state effects make accounting for the transfer free energies of all species significantly more challenging.

## II. EXPERIMENTAL METHODS

In the following sections we describe how we measured cyclohexane/water distribution coefficients for the 53 compounds displayed in Figure 1. The compound selection procedure is described in Section II A.

Distribution coefficient measurements utilized a shake-flask approach based on a liquid chromatography-tandem mass spectrometry (LC-MS/MS) technique previously developed for 1-octanol/water distribution coefficient measurements [21]. The approach is described in Section II B, and the procedure is schematically summarized in Figure 2.

**FIG. 1:**
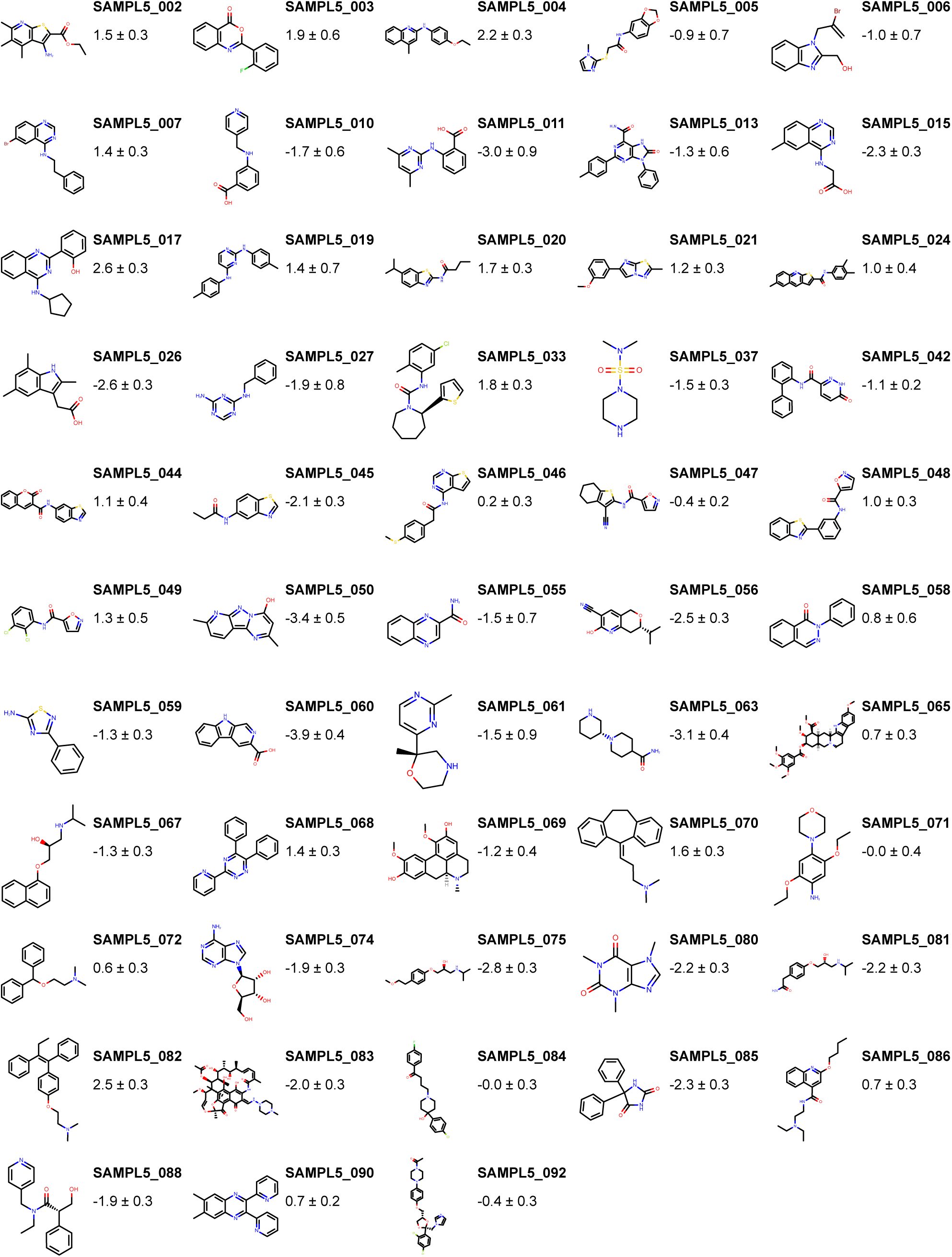
Molecules and corresponding measured log distribution coefficients for measurements that passed quality controls. Log D measurements are reported as expectation ± standard errors, calculated using our parametric bootstrap method (Section II D).

**FIG. 2:**
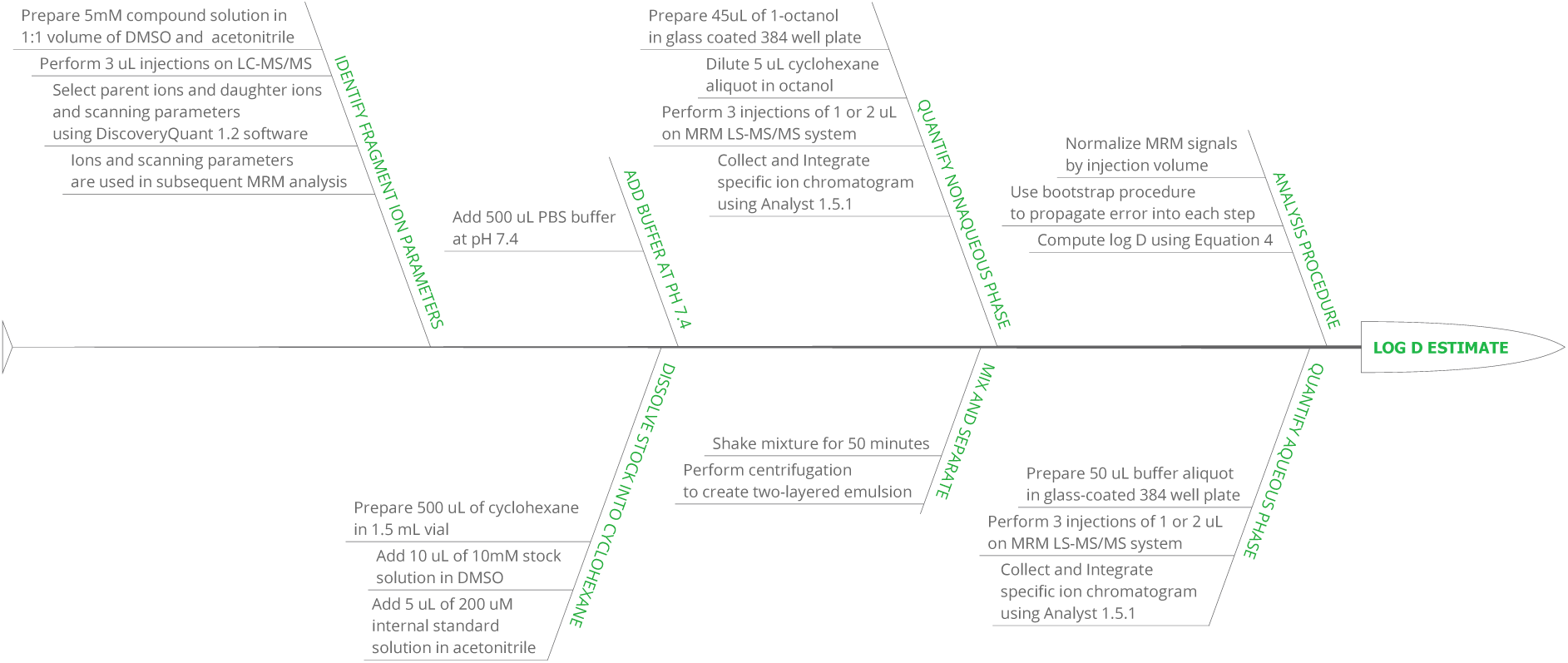
Illustration of the shake-flask procedure used for cyclohexane-water distribution coefficient measurements.

The measured data was subjected to a quality control procedure that eliminated measurements thought to be too unreliable for use in the SAMPLS challenge (Section IIC). Remaining data were analyzed using a physical model of the experiment by means of a parametric bootstrap procedure. We compared this approach to a nonparametric bootstrap approach, and the arithmetic mean and standard error of the data without bootstrap analysis. In Section II D, we describe each approach. The results for each approach can be found in Table I.

**Table I:** Log distribution coefficient measurements and standard errors. Estimates of log distribution functions and their associated standard errors are described for parametric bootstrap (Section II D 1), nonparametric bootstrap (Section II D 2), and arithmetic mean and corrected sample variance (Section II D 3).

| Compound ID | Uncertainty analysis method |  |  |
| --- | --- | --- | --- |
|  | Bootstrap |  | Arithmetic mean |
|  | Parametric | Nonparametric | Standard error |
| SAMPL5_002 | 1.5 ± 0.3 | 1.5 ± 0.2 | 1.4 ± 0.1 |
| SAMPL5_003 | 1.9 ± 0.6 | 1.9 ± 0.5 | 1.94 ± 0.04 |
| SAMPL5_004 | 2.2 ± 0.3 | 2.2 ± 0.1 | 2.2 ± 0.1 |
| SAMPL5_005 | -0.9 ± 0.7 | -0.9 ± 0.7 | -0.86 ± 0.03 |
| SAMPL5_006 | -1.0 ± 0.7 | -1.0 ± 0.6 | -1.02 ± 0.03 |
| SAMPL5_007 | 1.4 ± 0.3 | 1.39 ± 0.08 | 1.38 ± 0.09 |
| SAMPL5_010 | -1.7 ± 0.6 | -1.7 ± 0.5 | -1.7 ± 0.1 |
| SAMPL5_011 | -3.0 ± 0.9 | -3.0 ± 0.9 | -2.96 ± 0.03 |
| SAMPL5_013 | -1.3 ± 0.6 | -1.3 ± 0.5 | -1.5 ± 0.1 |
| SAMPL5_015 | -2.3 ± 0.3 | -2.3 ± 0.2 | -2.25 ± 0.09 |
| SAMPL5_017 | 2.6 ± 0.3 | 2.6 ± 0.2 | 2.5 ± 0.1 |
| SAMPL5_019 | 1.4 ± 0.7 | 1.4 ± 0.7 | 1.2 ± 0.1 |
| SAMPL5_020 | 1.7 ± 0.3 | 1.7 ± 0.1 | 1.6 ± 0.1 |
| SAMPL5_021 | 1.2 ± 0.3 | 1.18 ± 0.07 | 1.2 ± 0.1 |
| SAMPL5_024 | 1.0 ± 0.4 | 1.0 ± 0.4 | 1.0 ± 0.1 |
| SAMPL5_026 | -2.6 ± 0.3 | -2.6 ± 0.1 | -2.58 ± 0.04 |
| SAMPL5_027 | -1.9 ± 0.8 | -1.9 ± 0.7 | -1.87 ± 0.02 |
| SAMPL5_033 | 1.8 ± 0.3 | 1.82 ± 0.07 | 1.80 ± 0.08 |
| SAMPL5_037 | -1.5 ± 0.3 | -1.54 ± 0.07 | -1.53 ± 0.03 |
| SAMPL5_042 | -1.1 ± 0.2 | -1.13 ± 0.04 | -1.1 ± 0.1 |
| SAMPL5_044 | 1.1 ± 0.4 | 1.1 ± 0.3 | 1.0 ± 0.1 |
| SAMPL5_045 | -2.1 ± 0.3 | -2.09 ± 0.04 | -2.09 ± 0.08 |
| SAMPL5_046 | 0.2 ± 0.3 | 0.19 ± 0.09 | 0.20 ± 0.09 |
| SAMPL5_047 | -0.4 ± 0.2 | -0.37 ± 0.07 | -0.37 ± 0.09 |
| SAMPL5_048 | 1.0 ± 0.3 | 1.0 ± 0.2 | 0.9 ± 0.1 |
| SAMPL5_049 | 1.3 ± 0.5 | 1.3 ± 0.5 | 1.28 ± 0.04 |
| SAMPL5_050 | -3.4 ± 0.5 | -3.4 ± 0.5 | -3.2 ± 0.2 |
| SAMPL5_055 | -1.5 ± 0.7 | -1.5 ± 0.7 | -1.48 ± 0.04 |
| SAMPL5_056 | -2.5 ± 0.3 | -2.46 ± 0.05 | -2.46 ± 0.05 |
| SAMPL5_058 | 0.8 ± 0.6 | 0.8 ± 0.5 | 0.82 ± 0.03 |
| SAMPL5_059 | -1.3 ± 0.3 | -1.34 ± 0.03 | -1.33 ± 0.09 |
| SAMPL5_060 | -3.9 ± 0.4 | -3.9 ± 0.3 | -3.87 ± 0.08 |
| SAMPL5_061 | -1.5 ± 0.9 | -1.5 ± 0.9 | -1.45 ± 0.03 |
| SAMPL5_063 | -3.1 ± 0.4 | -3.2 ± 0.4 | -3.0 ± 0.1 |
| SAMPL5_065 | 0.7 ± 0.3 | 0.7 ± 0.1 | 0.69 ± 0.07 |
| SAMPL5_067 | -1.3 ± 0.3 | -1.3 ± 0.2 | -1.3 ± 0.1 |
| SAMPL5_068 | 1.4 ± 0.3 | 1.4 ± 0.2 | 1.41 ± 0.09 |
| SAMPL5_069 | -1.3 ± 0.4 | -1.2 ± 0.3 | -1.3 ± 0.1 |
| SAMPL5_070 | 1.6 ± 0.3 | 1.6 ± 0.2 | 1.61 ± 0.09 |
| SAMPL5_071 | -0.0 ± 0.4 | -0.0 ± 0.4 | -0.1 ± 0.2 |
| SAMPL5_072 | 0.6 ± 0.3 | 0.6 ± 0.2 | 0.6 ± 0.1 |
| SAMPL5_074 | -1.9 ± 0.3 | -1.9 ± 0.2 | -1.9 ± 0.1 |
| SAMPL5_075 | -2.8 ± 0.3 | -2.8 ± 0.1 | -2.77 ± 0.09 |
| SAMPL5_080 | -2.2 ± 0.3 | -2.2 ± 0.1 | -2.18 ± 0.07 |
| SAMPL5_081 | -2.2 ± 0.3 | -2.2 ± 0.1 | -2.19 ± 0.09 |
| SAMPL5_082 | 2.5 ± 0.3 | 2.5 ± 0.2 | 2.5 ± 0.1 |
| SAMPL5_083 | -2.0 ± 0.3 | -2.0 ± 0.2 | -1.9 ± 0.1 |
| SAMPL5_084 | -0.0 ± 0.3 | -0.02 ± 0.05 | -0.02 ± 0.08 |
| SAMPL5_085 | -2.3 ± 0.3 | -2.3 ± 0.2 | -2.2 ± 0.1 |
| SAMPL5_086 | 0.7 ± 0.3 | 0.7 ± 0.1 | 0.70 ± 0.06 |
| SAMPL5_088 | -1.9 ± 0.3 | -1.9 ± 0.2 | -1.9 ± 0.1 |
| SAMPL5_090 | 0.7 ± 0.2 | 0.75 ± 0.06 | 0.76 ± 0.08 |
| SAMPL5_092 | -0.4 ± 0.3 | -0.41 ± 0.09 | -0.39 ± 0.09 |

### A. Compound selection

Compounds were initially selected from a database of is 9115 lead-like molecules available in eMolecules that were iss present in the Genentech chemical stores in quantities of over 2 mg, with molecular weights between 150-350 Da. The lower bound on molecular weight was chosen to increase the likelihood of detectability by mass spectrometry, and the upper bound to limit molecular complexity.

We initially chose approximately 88 compounds based on several criteria:

- First, we selected 8 carboxylic acid compounds. These were of potential interest for the purpose of the challenge, since it was suspected these could potentially partition along into the cyclohexane phase together with water or cations [23].
- The software MoKa, version 2.5 was used to obtain calculated LogP, LogD, and pKa values [24, 25]. This version of MoKa was trained with Roche internal data to improve accuracy. We selected 20 compounds with predicted pKa values that would potentially be measurable with a Sirius T3 instrument (Sirius Analytical) so validation with an orthogonal technique (electrochemical titration) could be performed in the future. The pKa predictions for compounds in our final data set have been made available in the Supplementary Information.
- The remaining compounds were divided into 10 equal-size bins that spanned the predicted dynamic range of log P values (-3.0 to 6.6), and 6 compounds were drawn from each bin, to a total of 60.

This set of 88 molecules was later reduced to 64 molecules due to the unavailability of some compounds or the inability to detect molecular fragments by mass spectrometry at the time of measurement. This selection was expanded to include 31 compounds used as internal standards for the previously developed octanol/water assay protocol [21], bringing the total number of compounds for which measurements were performed to 95. These compounds were randomly assigned numerical SAMPL_XXX designations for the SAMPL5 blind challenge. After the quality control filtering phase (Section II C), the resulting data set contained 53 compounds, which are displayed in Figure 1. Canonical isomeric SMILES representations for the compounds can also be found in Table S1. These were generated using OpenEye Toolkits v2015.June by converting 3D SDF files, after manually verifying the correct stereochemistry.

### B. Shake-flask measurement protocol for cyclohexane/water distribution coefficients

We adapted a shake-flask assay method from an original octanol/water LC-MS/MS protocol [21] to accommodate the use of cyclohexane for the nonaqueous phase. Our modified protocol is described here, and the procedure is explained schematically in Figure 2.

The log D is estimated by quantifying the concentration of a solute directly from two immiscible layers, present as an emulsion in a single vial. Capped glass 1.5 mL auto-injector vials with PTFE-coated silicone septa^1^ were used for partitioning, as cyclohexane was found to dissolve polystyrene 96-well plates used in the original protocol.

For each individual experiment, 10µL of 10 mM compound in dimethyl sulfoxide (DMSO)^2^ and 5 µL of 200 µM propanolol in acetonitrile (an internal standard) were added to 500 µL cyclohexane^3^, followed by the addition of 500 µL of PBS solution^4^. The ionic components of the buffer were chosen to replicate the buffer conditions used in other in-vitro assays at Genentech. Unlike the original protocol, neither phase was presaturated prior to pipetting.

The solute was allowed to partition between solvents while the mixture was shaken for 50 minutes using a plate shaker^5^ at 800 RPM, while the vials were mounted in a vial holder and taped down to the sides of the vial holder^6^. The two solvents were then separated by centrifugation for 5 minutes at 3700 RPM in a plate centrifuge, using the plate rotor^7^, with the vials seated in the same vial holder.

Aliquots were extracted from each separated phase using a standard adjustable micropipette, and transferred into a 384-well glass-coated polypropylene plate for subsequent quantification^8^. Cylcohexane wells were first prepared with 45 µL of 1-octanol^9^ per well. 5 µL of cyclohexane was extracted from the top phase by micropipette and mixed with 45 µL of octanol in the 384 well plate. 50 µL of aqueous solution was subsequently extracted from the bottom phase. The octanol dilution was performed mainly to prevent accumulation of cyclohexane on the C18 HPLC columns^10^ that were used. For the aqueous (bottom) phase, the aliquot of 50 µL was transferred directly into the 384-well plate, into wells that did not contain octanol. The 384-well plates were sealed with using glueless aluminum foil seals^11^, and fragment concentrations assayed using quantitative LC-MS/MS.

Measuring solute distribution into the two phases depends on two separate mass spectrometry measurements^12^:

- The solute is analyzed to identify and select parent and daughter ions, and optimize ion fragment parameters^13^. We used a flow rate of 0.2 mL/min, mobile phase of water/acetonitrile/formic acid (50/50/0.1 v/v/v) and 1.5 min run time. All parameters were automatically stored for further multiple-reaction monitoring (MRM) analysis. For several compounds, the fragment identification LC-MS/MS procedure did not yield high intensity fragments, and these could therefore not be measured using the MRM approach. All identified parent and daughter ions are available as part of the Supplementary Information.
- A separate mass spectrometer is employed using MRM to select for parent ions and daughter ions of the solute identified in the previous step. The mass/charge (m/z) intensity (proportional to the absolute number of molecules) is quantified as a function of the retention time^14^. Information on the gradient can be found in Supplementary Table 1 of Lin and Pease 2013 [21].

Highest m/z intensity fragments were selected using 5 mM solutions consisting of 50% DMSO, 50% acetonitrile.

From each solvent phase in the partitioning experiment, one aliquot was prepared, and replicate MRM measurements were performed 3 times per aliquot. The log D can be calculated from the relative MRM-signals, obtained by integrating the single peak in the MRM-chromatogram, using Equation (4).

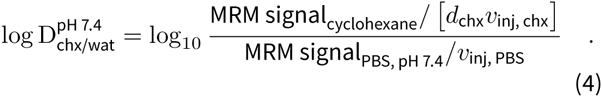

The cyclohexane signal is normalized by the dilution factor of our cyclohexane aliquots, *d*_chx_ = 0.1, and the injection volume *v*_inj, chx_. As the PBS aliquots were not diluted, this is only normalized by the injection volume *v*_inj,PBS_. Experiments were carried out independently at least in duplicate, repeated from the same DMSO stock solutions. Injection volumes of the MRM procedure were 1 µL for cyclohexane (diluted in octanol), and 2 µL for PBS samples. To optimize experimental parameters, we carried out two additional repeat experiments with 2 µL injections for cyclohexane (diluted in octanol), and 1 µL for PBS. This set included SAMPL5_003, SAMPL5_005, SAMPL5_006, SAMPL5_011, SAMPL5_027, SAMPL5_049, SAMPL5_050, SAMPL5_055, SAMPL5_058, SAMPL5_060, and SAMPL5_061. The additional repeats were carried with higher cyclohexane injection volumes to increase the strength of signal in the cyclohexane phase, and lower PBS volumes to decrease the chances of oversaturation of PBS phase signals.

### C. Quality control

In order to eliminate measurements thought to be too unreliable for the SAMPL5 challenge, we utilized a simple quality control filter after MRM quantification. Compounds where the integrated MRM signal within either phase varied between replicates or repeats by more than a factor of 10 were excluded from further analysis. We additionally removed compounds that exceeded the dynamic range of the assay because they did not produce a detectable MRM signal in either the cyclohexane or buffer phases during the quantification.

### D. Bootstrap analysis

Since our ultimate goal is to compare predicted distribution coefficients to experiment to evaluate the accuracy of current-generation physical modeling approaches, it is critical to have an accurate assessment of the uncertainty in the experimental measurement. Good approaches to uncertainty analysis propagate all known sources of experimental error into the final estimates of uncertainty. To accomplish this, we developed a parametric bootstrap model [26] of the experiment based on earlier work [27], with the goal of propagating pipetting volume and technical replicate errors through the complex analysis procedure to estimate their impact on the overall estimated log D measurements.

Bootstrap approaches provide new synthetic data sets, denoted as realizations, sampled using some function of the observed data that approximates the distribution that the observed data was drawn from. For each compound that was measured, suppose our data set provides N independent repeats (from the same stock solution, typically 2 or 4), and 3 technical replicates for each repeat (quantitation experiments from each repeat, typically 3). Each realization of the bootstrap process leads to a new synthetic data set, of the same size, from which a set of synthetic distribution coefficients can be computed for the realization. We applied two additional approaches for comparison to assess the performance of our parametric bootstrap method (Section II D1). One features a nonparametric bootstrap approach (Section II D2), which does not include any physical details. The other is a calculation of the arithmetic mean and standard error that is limited to the observed data (Section II D 3).

#### 1. parameteric bootstrap

We used a parametric bootstrap [28] method to introduce a random bias and variance into the data, based on known experimental sources. This procedure allows us to use a model to propagate known uncertainty throughout the procedure [28]. This allows us to better estimate the distribution that the observed data was drawn from, so that more accurate estimates of the means and sample variance can be obtained.

Uncertainties in pipetting operations were modeled based on manufacturer descriptions [29, 30], following the work of Hanson, Ekins and Chodera [27]. Technical replicate variation was modeled by calculating the coefficient of variation (CV) between individual experimental replicates. We then took the mean CV of the entire data set, which was found to be ~0.3. As a control, we verified that the CV did not depend on the solvent phase that was measured. We included this in the parametric model by adding a signal imprecision, modeled by a normal distribution with zero mean, and a standard deviation of 0.3. We perform a total of 5 000 realizations of this process, and calculate statistics overall realizations, such as the mean (expectation) and standard deviation (estimate of standard error) for each measurement.

#### 2. Nonparameteric bootstrap

A traditional nonparametric Monte Carlo procedure was applied to resample data points[26]. This approach can estimate the distribution that the observed data was drawn from by resampling from the observed data with replacement, to generate a new set of data points with size equal to the observed data set. Nonparametric bootstrap can be a useful approach if larger amounts of data are available, and a detailed physical model of the experiment is absent. We implemented the procedure in two stages:

1. A set of *N* repeats is drawn with replacement from the original set of measured repeats.
2. For each of the repeats, we similarly draw a set of 3 technical replicates from the original set of technical replicates.

This yields a sample data set with the same size as the originally observed data (*N* repeats, with 3 replicates each). We perform a total of 5 000 realizations of this process, and calculate statistics over all realizations, such as the mean (expectation) and standard deviation (estimate of standard error) for each measurement.

#### 3. Arithmetic mean and sample variance

We calculated the arithmetic mean over all replicates and repeats, and estimated the standard error from the total of 6 or 12 data points, to compare to our bootstrap estimates.^15^

### E. Kernel densities

As a visual guide, in Figure 3 data are plotted on top of an estimated density of points. This density was calculated using kernel density estimation [31], which is a nonparametric way to estimate a distribution of points using kernel functions. Kernel functions assign density to individual points in a data set, so that the combined set of data points reflects a distribution of of the data. We used the implementation available in the python package seaborn, version 0.7.0 [32]. We used a product of Gaussian kernels, with a bandwidth of 0.4 for log D and 0.3 for the standard error. To prevent artifacts such as negative density estimates for the standard errors, they were first transformed by the natural logarithm ln, and the results were then converted back into standard errors by exponentiation.

**FIG. 3:**
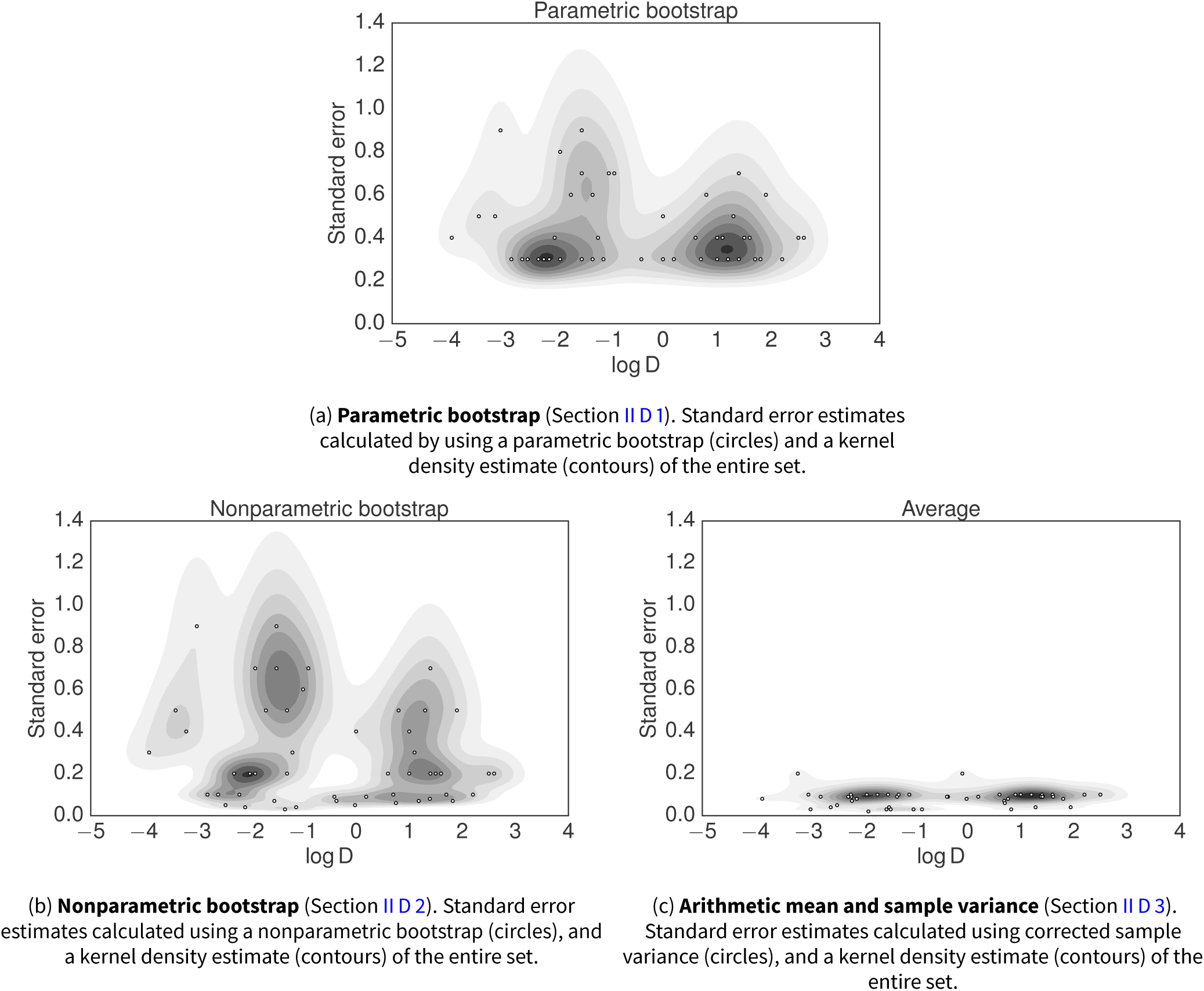
Joint kernel density estimates of log distribution coefficient (log D) measurements and measurement error estimates. `log` D measurements are plotted with their corresponding estimated standard errors (circles) for the three analysis approaches described in Section II D. A kernel density estimate (contours, described in Section II E) is shown to highlight the differences in error estimates for the different methods.

## III. DISTRIBUTION COEFFICIENTS

The log D values and their uncertainties for the 53 small molecules that passed quality controls are presented in Table I. In the following two sections, we describe the differences between the analysis results in more detail.

### A. Mean and standard errors in log D

The results from the arithmetic mean and sample variance calculation ( Section II D 3) are plotted in Figure 3c. Despite the compound selection effort, the distribution of data along the log D-axis is less dense in the region -1 to 0 log units. The data outside this region seems to be centered around -2 log units, or around 1 log unit. We could attribute this distribution of data to coincidence, though this may warrant future investigations into systematic errors. Using the arithmetic mean of the combined repeat and replicate measurements (Section II D 3) the distribution coefficients measured spanned from -3.9 to 2.5 log units.

The log D measurements distribution appears bimodal along the uncertainty axis. A subset of mostly negative log D values (Figure 3c) has a smaller estimated standard deviation, though this is not the case for the majority of negative log D values. The average standard error, rounded to 1 significant figure, is 0.2 log units for the arithmetic mean calculation.

### B. Bootstrap results

Estimates of the log D span the range between -3.9 to 2.6 log units, using either of the two bootstrap approaches (Section II D 1 and Section II D 2). The log D estimates do not differ significantly from the arithmetic mean calculations. The difference between the results is seen when we compare the estimated standard errors. When applying our bootstrap procedures (Section II D 1 and Section II D 2), we see an upwards shift in the uncertainties, compared to the sample variance calculations. The nonparametric approach yields an average uncertainty of 0.3 log units. The parametric approach yields an average uncertainty of 0.4 log units. The parametric bootstrap suggests that by propagating errors such as the cyclohexane dilution, and the replicate variability into the model, some of the observed low uncertainties might be an artifact of the low number of measurements. This suggest that simply calculating the arithmetic mean, and standard error of all measured data might not reliably capture the error in the experiments. We also note that for certain compounds, bootstrap distributions exhibit multimodal character and as such, standard errors might not accurately capture the full extent of the experimental uncertainty. We provide the bootstrap sample distributions of the parametric model in the supplementary information.

Using the parametric scheme, we see an average shift of uncertainties to larger values compared to the nonparametric bootstrap. The density estimate suggests we should expect a lower bound to the error that we have now incorporated into the analysis. Not every compound shows the same increase in uncertainty, though if we compare the two bootstrap approaches, results are similar above this empirically observed lower bound. The nonparametric approach returns higher uncertainties for some data on average, but estimates lower uncertainties for some as well. It can be concluded that the error would typically be underestimated without the use of a bootstrap approach. Without a physical model, a nonparametric approach might still underestimate errors due to the limited sample size for each measurement (either 2 or 4 fully independent repeats, and a total of 3 replicates per data point).

### C. Correlation of uncertainty with physical properties

We investigated whether there was an obvious correlation between the uncertainty estimates obtained from our analysis and the properties of the molecules in our data set. A set of simple physical descriptors including molecular weight, predicted net charge, and the total number of amines and hydroxyl moieties were plotted against the bootstrap uncertainty. None of the descriptors tested had an absolute Pearson correlation coefficient R whose 95% confidence interval did not contain the correlation-free R = 0, according to methods described by Nicholls [33]. The analysis can be found in the Supplementary Information as an Excel spreadsheet.

## IV. DISCUSSION

### 1. Solvent conditions

It is important to consider the influence of cosolvents on the measured values. The solutions contained approximately 1% DMSO, as well as approximately 0.5% acetonitrile. Further work would benefit from a comparison with experiments starting from dry stocks, and thereby not adding extra solvents. This would eliminate DMSO and acetonitrile, by dispensing compound directly into either cyclohexane, or the mixture of cyclohexane and PBS. In this case, care must be taken that the compound is fully dissolved. If found to be necessary all experiments could be started from dry compound stocks, to entirely eliminate effects from cosolvents such as DMSO and acetonitrile. This would make experiments more laborious, and would therefore reduce the bandwidth of the method.

Differential evaporation rates of cyclohexane and water could be an additional source of error. Cyclohexane (vapor pressure 97.81 torr [34]) is more volatile than water (vapor pressure 23.8 torr[35]). Evaporation from the cyclohexane-water phase-separated mixture or aliquots from individual phases could increase the concentration of the cyclohexane phase more rapidly than the water phase, leading to an over-estimation of the log D. For future investigations, it would be prudent to verify that evaporation rates are sufficiently low to ensure no significant impact on the measured log D.

### 2. Compound detection limits

Calculations using COSMO-RS software[36] suggested a systematic underestimation of log D values in the negative log unit range, in particularly past a log D of -2. Without further experimental investigation, we can not draw definite conclusions as to whether this is the case, or if so, where the source of the systematic error lies.

One possibility that may cause an artificial reduction of the dynamic range—especially at high log D values—is the potential for MS/MS detector saturation at high ligand concentrations. Previous work (Figure 2 from [21]) examined detector saturation effects, finding it possible to reach sufficiently high compound concentrations (generally ≥10 µM) that MRM is no longer linear in compound concentration for that phase. This work also found that different compounds reach detector saturation at different concentrations [21], in principle requiring an assessment of detector saturation to be performed for each compound. While we could not deduce obvious signs of detector saturation in our LC-MS/MS chromatograms, these effects could be mitigated by performing a dilution series of the aliquots sampled from each phase of the partitioning experiment to ensure detector response is linear in the range of dilutions measured. This may also reveal whether compound dimerization may be a complicating factor in quantitation.

### 3. Experimental design considerations

In order to adjust our experimental setup, we had to switch away from using polystyrene 96 well plates, as these were dissolved by cyclohexane. We attempted the use of glass inserts, and glass tubes but these were too narrow and provided insufficient mixing when shaken. We switched to glass vials because their larger diameter provides improved mixing when shaken. For future work, we would recommend the use of glass coated plates, which have the automation advantages of the plates used in the original protocol [21].

Plate seals need to be selected carefully. We experimented with silicone sealing mats, but these absorbed significant quantities of cyclohexane. We also had to discontinue use of aluminum seals that contained glue, since the glue is soluble in cyclohexane and would contaminate LC-MS/MS measurements. In the end, we used aluminum PlateLoc heat seals and glass coated 384 well plates to circumvent these issues.

Sensitivity also suffered due to the need to dilute cyclohexane in octanol to prevent its accumulation on C18 columns used in the LC-MS/MS phase of the experiment. Trial injections on a separate system and chromatograms showed accumulation of unknown origin at the end of each UV chromatogram. Accumulation was reduced by injecting less cyclohexane. As a result, we diluted the cyclohexane with 1-octanol for the experiments described here, and ran blank injections containing ethanol between batches of 64 measurements to ensure the column was clean.

Another change to the protocol that we would like to consider for future measurements is to optimize the time spent equilibrating the mixture. In this work, we separated phases via centrifugation and sampled aliquots for concentration measurement within minutes. The post-centrifugation time prior to sampling aliquots could be extended to 24 hours to allow for more equilibration for the solute between phases. This may have a downside, since we would have to consider the effects that may follow if compounds prefer to be in the interface-region between cyclohexane and water, or water and air. These could cause high local concentrations, introducing a dependency of the results on exactly which part of the solution aliquots are taken from. We can get around this by only taking samples from the pure cyclohexane and aqueous regions, avoiding the interfaces. This way, we still get the right distribution coefficients for partitioning between bulk phases even if some compound is lost to the interfaces.

It may be worthwhile to consider other effects of pipetting operations on the procedure. Some compounds could potentially stick to the surface of pipettes, or glass surfaces. This could adversely affect our measurements by changing local concentrations.

We also consider that assay results might be less variable if we presaturated water and cyclohexane before mixing them. While cyclohexane and water have much lower mutual solubility than octanol, it is still possible that this affects the measurement.

For future challenges, we would recommend that these assays are carried out at multiple final concentrations of the ligand in the assay. This could be achieved using different volumes of 10 mM ligand stocks. This would help detect dimerization issues, and may help account for issues with detector oversaturation. Note that the absolute errors in these stock volumes will not be critical, since the measurements rely on the relative measurement between the two phases. We could build models that allow for extrapolation to the infinite dilution limit, which should then provide simpler test cases for challenge participants to reproduce. On the opposite end, it may be useful to even investigate ways to design an experimental set that represents these type of issues, such as compound dimerization, so that we can focus more on these.

### 4. Uncertainty analysis

We hope the experience from this challenges will lay the groundwork for improving the reliability of data sets regarding the physical properties that we as a modeling community rely on. Many computational studies are limited in the amount of high-quality experimental data that they have access to. Unfortunately, most data are taken straight from literature tables, without much thought being spent on the data collection process. By performing the experimental part of the SAMPL5 challenge we were in the position to provide new data to the modeling community, with an opportunity to decide on an analysis strategy that suits modeling applications. This not only allows for blind validation of physics-based models, but also a re-evaluation of the exact properties a data set should have to provide utility to the modeling community. An important fact that we feel needs reemphasizing is that experimental data are limited in utility by the method that was used to analyze it.

Among the lessons learned from this challenge, we would recommend that future challenges would also feature a rigorous statistical treatment of the experimental analysis procedure, ideally going beyond these initial efforts. One crucial part of the analysis procedure is obtaining not only accurate estimates of the observable, but also its uncertainty. As indicated in our data set, standard error estimates from small populations may underestimate the error. Several approaches can be taken to resolve part of this issue. Among the options are the use of statistical tests, such as the bootstrapping methods we applied in this work. These can help us both propagate information on uncertainty into the model (such as a parametric bootstrap) or extract uncertainty already available in the data (such as nonparametric bootstrap). The parametric approaches can be improved in terms of the physical models that are used to analyze the data. These models should ideally include all known sources of error, such as pipetting errors, evaporation of solvent, errors in integration software, fluctuations in temperature, pressure and likewise many other conditions that could affect the results.

Another approach would be to perform statistical inference on the data set, to provide uncertainty estimates from the data itself. The model structure can provide ways to incorporate data and propagate uncertainty from multiple experiments. Common parameters, such as variance in measurements between experiments could be inferred from combining the entire data set into one model. When prior knowledge on the experimental parameters is available, a Bayesian model can be used to effectively infer this type of uncertainty from the data, and use it to propagate the error into log D estimates. Distinctions could be made between an objective treatment of the problem, or an empirical Bayesian approach, where prior parameters are derived from the data. One could use a maximum *aposteriori*(MAP) probability approach to obtain an estimate of one of the modes of the parameter distribution. This has obvious downsides when posterior densities are multimodal, and in such a case, one may wish to estimate the shape of the entire posterior distribution instead. An approach like Markov chain Monte Carlo (MCMC) [37] could provide such estimates, and will allow for calculation of credible intervals. MCMC methods can be computationally intensive compared to MAP, though if the resulting posterior is complicated, a MAP estimate can give poor results. Unfortunately, we were unable to construct a Bayesian model of the experiments within our time constraints. We would encourage future challenges to make an attempt at creating a Bayesian model, since this would allow for robust inference of all experimental parameters.

### 5. Funding future challenges

The execution of this work would not have been possible without the resources provided by Genentech. Access to a rich library of compounds onsite allowed us to select a dataset that was both challenging and useful for the purposes of the SAMPL challenge. At the same time, the instrumentation provided us with the bandwidth to perform many measurements. Rapid redesign of experiments by trial and error, as a result of the difficulties with cyclohexane compatibility of laboratory consumables and equipment, would not have been possible without the expertise shared by Genentech scientists and the opportunities to do many measurements.

Future iterations of this challenge would benefit from continued collaboration between industry and academia. Academic groups can partner with industry groups to pair available skilled academic labor (graduate students and postdoctoral researchers) with specialized measurement equipment and compound libraries. The graduate student industry internship model proved to be a particularly successful approach, with measurements for a blind challenge providing a well-defined, limited-scope project with clear high value to the modeling community.

## V. CONCLUSION

The experimental data provided by this study was very useful for hosting the first small-molecule distribution coefficient challenge in the context of SAMPL. It revealed that log D prediction, as well as measurement, is not always straightforward. We showed that it was possible to perform cyclohexane/water log D measurements in the same manner as the original octanol/water assays, though further optimizations are needed to reach the same level of throughput. Cyclohexane did pose several challenges for experimental design, such as the need for different container types, and the potential accumulation of substrate on reversed phase HPLC columns.

Many details, such as protonation states, tautomer states, and dimerization might need to be accounted for in order to reproduce experiments. This challenge taught us considerations that should be made on the experimental side. Cases where dimerization were pointed out as possible reason for discrepancy between experiment and model, could only be hypothesized from the modeling end and not tested experimentally. Issues with detector saturation could also be affecting the overall quality of the data set. Future experiments would benefit from more rigorous protocols, such as measurements at multiple concentrations, and models of all experimental components.

We recommend that future challenges, and experiments in general, use physical models of experiments in the analysis of experimental uncertainty. These should be part of the analysis procedure, but also in experimental design. These will reveal abnormalities in data more clearly.

We recommend that future challenges look into the use of bootstrap models such as those considered here. Additionally, the use of Bayesian inference methods, that allow the incorporation of prior information should lead to a more robust estimate of experimental uncertainty. They will allow for joint inference on multiple experiments, thereby increasing the information gain by using the model.

Lastly, the sponsoring of this internship by Genentech was fundamental to generating this data. Access to compound libraries, and the equipment to perform the experiments is crucial to the design and execution of a study. Close collaborations with Genentech scientists were important in solving many technical challenges. The collaboration between industry and academics was not only fruitful, but fundamental in establishing standardized challenges for the modeling field. The amount of data we were able to gather would have been hard to come by without industry resources. At the same time, the need and expertise in investigating these challenging physical chemical problems provided by the community, and the forum provided by the SAMPL challenge was essential in turning this challenge into a success. We welcome such future efforts and collaborations, as it is apparent that both experimental and computational approaches for obtaining log D estimates for small molecules, would benefit from further optimization.

## VI. SUPPLEMENTARY INFORMATION

Canonical isomeric smiles for each of the measured compound are available in Table S1. An sdf file containing all compounds, including the measured distribution coefficients is available as part of the Supplementary Information. Parent and daughter fragment ion information is available as part of the Supplementary Information. Integrated MRM data including excluded data points are available as part of the Supplementary Information. Bootstrap distributions from the parametric bootstrap samples for each compound are provided in the Supplementary Information. A correlation analysis between the parametric bootstrap uncertainty, and the chemical properties of the compounds in the dataset is available as an Excel spreadsheet in the Supplementary Information. We also include a csv file containing a full list of SAMPL5_XXX identifiers and canonical isomeric smiles, including unmeasured compounds. Source code of the bootstrap uncertainty analysis is available on Github at https://github.com/choderalab/sampl5-experimental-logd-data. A copy of this source code is also included in a zip file, as part of the supporting information.

## IX. ACKNOWLEDGMENTS

The authors acknowledge Christopher Bayly (OpenEye Scientific) and Robert Abel (Schrödinger) for their contributions to discussions on compound selection; Joseph Pease (Genentech) for discussions of the experimental approach and aid in compound selection; Delia Li (Genentech) for her assistance in performing experimental work; Alberto Gobbi (Genentech), Man-Ling Lee (Genentech), and Ignacio Aliagas (Genentech) for helpful feedback on experimental issues; Andreas Klamt (Cosmologic) and Jens Reinisch (Cosmologic) for invigorating discussions regarding experimental data; Patrick Grinaway (MSKCC) for helpful discussions on analysis procedures; and Anthony Nicholls (OpenEye) for originating and supporting earlier iterations of SAMPL challenges.

## X. SUPPLEMENTARY INFORMATION

### A. Compound identifiers

**TABLE S1:**
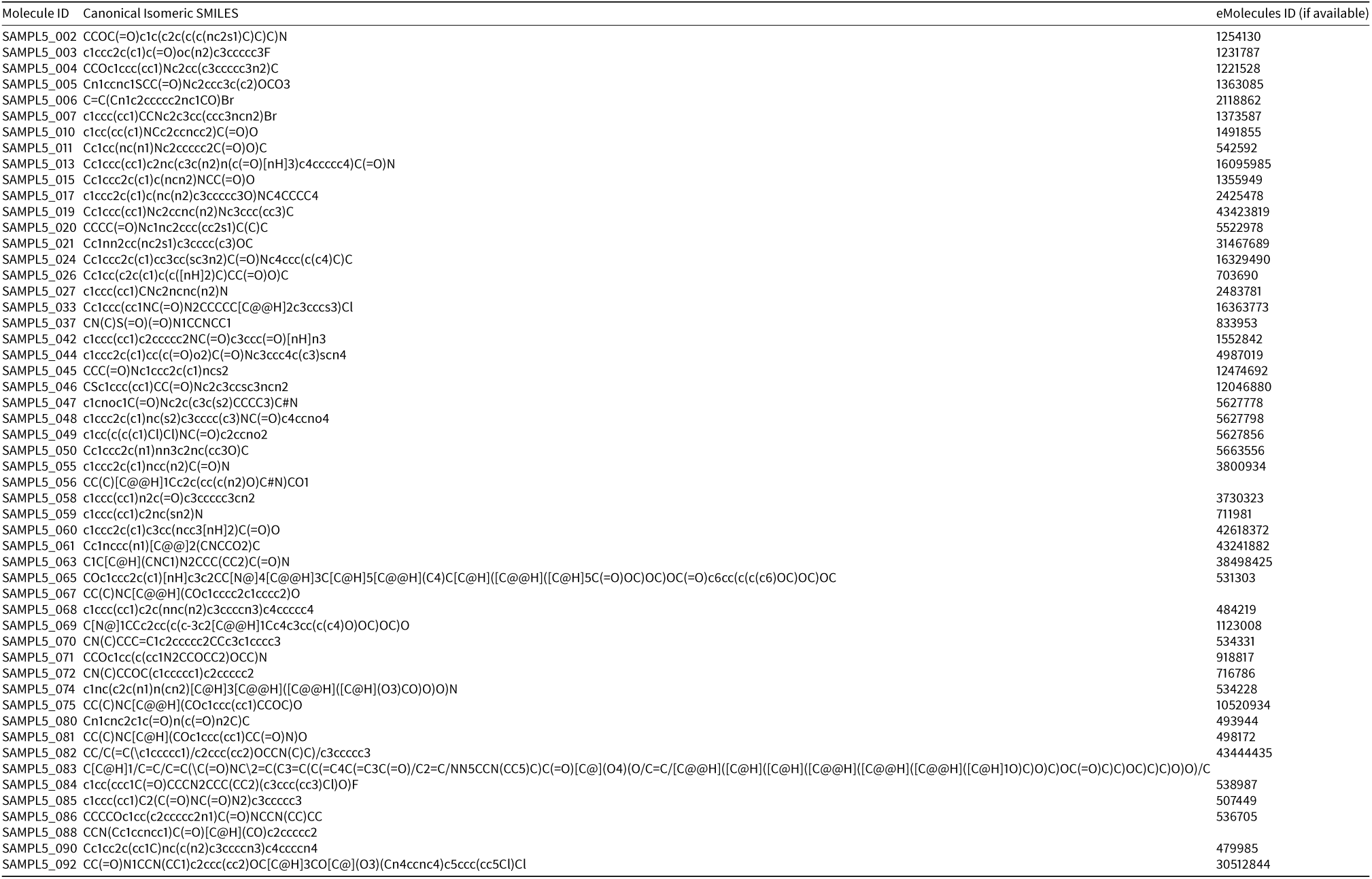
All of the compounds that were selected, and for which log D was obtained.

## Footnotes

1 Shimadzu cat. no. 22s-4s4s0-91

2 DMSO stocks from Genentech compound library

3 ACS grade ≥99%, Sigma-Aldrich cat. no i79i9i-2L, batch #00sssME

4 136 mM NaCl, 2.6 mM KCl, 7.96mMNaHPO,i.46mM KHPO, with pH adjusted to 7.4, prepared by the Genentech Media lab

5 Thermo Fisher Scientific, Titer Plate Shaker, model: 4625,Waltham, MA, USA

6 Agilent Technologies, Vial plate for holding 54 × 2 mL vials part no. G2255-68700

7 Eppendorf, Centrifuge 5804, Hamburg, Germany

8 384-well glass coat plate:Thermo Scientific, Microplate, 384-Well; Webseal Plate; Glass-coated Polypropylene; Square well shape; U-Shape well bottom; 384 wells; 90uL sample volume; catalog number: 3252187

9 ACROS Organics, 1-octanol99%pure, catalog number: AC150630010, Geel, Belgium

10 Waters Xbridge C18 2.130mmwith 2.5mparticles

11 Agilent cat no 24214-001

12 All LC solventswere HPLC-grade and purchased fromOmniSolv (Charlotte, NC, USA)

13 This was done using a Shimadzu NexeraX2 consisting of an LC-30AD(pump), SIL-30AC (auto-injector), and SPD-20AC(UV/VIS detector) with Sciex API4000QTRP (MS)

14 This was done using a Shimadzu NexeraX2 consisting of an LC-30AD(pump), SIL-30AC (auto-injector), and SPD-20AC(UV/VIS detector) with Sciex API4000 (MS)

15 For the purpose of the D3R/SAMPL5 workshop, we originally erroneously reported the standard deviation 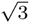 instead of the standard error 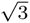. The factor of 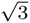 corrects the sample standard deviation across all MRM measurements for the correlation between the 3 replicate measurements belonging to a single independent experimental repeat.

## References

[1] J. P. Guthrie, J Phys Chem B 113, 4501 (2009).

[2] M. T. Geballe, A. G. Skillman, A. Nicholls, J. P. Guthrie, and P. J. Taylor, J Comput Aided Mol Des 24, 259 (2010).

[3] A. G. Skillman, J Comput Aided Mol Des 26, 473 (2012).

[4] H. S. Muddana, A. T. Fenley, D. L. Mobley, and M. K. Gilson, J Comput Aided Mol Des 28, 305 (2014).

[5] D. L. Mobley, K. L. Wymer, N. M. Lim, and J. P. Guthrie, J Comput Aided Mol Des 28, 135 (2014).

[6] P. Czodrowski, C. A. Sotriffer, and G. Klebe, J. Mol. Biol. 367, 1347 (2007).

[7] H. Steuber, P. Czodrowski, C. A. Sotriffer, and G. Klebe, J. Mol. Biol. 373, 1305 (2007).

[8] Y. C. Martin, J. Comput. Aid. Mol. Des. 23, 693 (2009).

[9] R. Mannhold, G. I. Poda, C. Ostermann, and I. V. Tetko, J. Pharm. Sci. 98, 861 (2009).

[10] P. A. Kollman, Accounts of Chemical Research 29, 461 (1996).

[11] S. A. Best, K. M. Merz, and C. H. Reynolds, The Journal of Physical Chemistry B 103, 714 (1999).

[12] B. Chen and J. I. Siepmann, The Journal of Physical Chemistry B 110, 3555 (2006).

[13] A. P. Lyubartsev, S. P. Jacobsson, G. Sundholm, and A. Laaksonen, J. Phys. Chem. B 105, 7775 (2001).

[14] N. Bhatnagar, G. Kamath, I. Chelst, and J. J. Potoff, J. Pharm. Sci. 137, 014502 (2012).

[15] S. A. Margolis and M. Levenson, Fresenius’ J. Anal. Chem. 367, 1 (2000).

[16] R. Stephenson, J. Stuart, and M. Tabak, Journal of Chemical & Engineering Data 29, 287 (1984).

[17] C. Black, G. G. Joris, and H. S. Taylor, The Journal of Chemical Physics 16, 537 (1948).

[18] S. H. Yalkowsky, Y. He, and P. Jain, Handbook of aqueous solubility data (CRC press, ADDRESS, 2010).

[19] J. G. Harris and F. H. Stillinger, J. Chem. Phys. 95, 5953 (1991).

[20] C. C. Bannan, G. Calabro, D. Y. Kyu, and D. L. Mobley, Journal of Chemical Theory and Computation (2016).

[21] B. Lin and J. H. Pease, Comb Chem High Throughput Screen 16, 817 (2013).

[22] W. M. Haynes, CRC handbook of chemistry and physics (CRC press, ADDRESS, 2014).

[23] A. Leo, C. Hansch, and D. Elkins, Chem. Rev. 71, 525 (1971).

[24] F. Milletti, L. Storchi, G. Sforna, and G. Cruciani, Journal of chemical information and modeling 47, 2172 (2007).

[25] F. Milletti, L. Storchi, L. Goracci, S. Bendels, B. Wagner, M. Kansy, and G. Cruciani, European journal of medicinal chemistry 45, 4270 (2010).

[26] B. Efron, Ann. Statist. 7, 1 (1979).

[27] S. M. Hanson, S. Ekins, and J. D. Chodera, Journal of computer-aided molecular design 29, 1073 (2015).

[28] B. Efron and R. J. Tibshirani, An introduction to the bootstrap (CRC press, ADDRESS, 1994).

[29] Rainin Pipet-Lite Multi Pipette L8-200XLS+, https://www.shoprainin.com/Pipettes/Multichannel-Manual-Pipettes/Pipet-Lite-XLS%2B/Pipet-Lite-Multi-Pipette-L8-200XLS%2B/p/17013805, accessed: 2016-06-06.

[30] Rainin Classic Pipette PR-10, https://www.shoprainin.com/Pipettes/Single-Channel-Manual-Pipettes/RAININ-Classic/Rainin-Classic-Pipette-PR-10/p/17008649, accessed: 2016-06-06

[31] M. Rosenblatt, Ann. Math. Statist. 27, 832 (1956).

[32] M. W. O. B. drewokane; Paul Hobson; Yaroslav Halchenko; Saulius Lukauskas; Jordi Warmenhoven; John B. Cole; Stephan Hoyer; Jake Vanderplas; gkunter; Santi Villalba; Eric Quintero; Marcel Martin; Alistair Miles; Kyle Meyer; Tom Augspurger; Tal Yarkoni; Pete Bachant; Constantine Evans; Clark Fitzgerald; Tamas Nagy; Erik Ziegler; Tobias Megies; Daniel Wehner; Samuel St-Jean; Luis Pedro Coelho; Gregory Hitz; Antony Lee; Luc Rocher;, seaborn: v0.7.0 (January 2016), 2016.

[33] A. Nicholls, Journal of Computer-Aided Molecular Design 28, 887 (2014).

[34] Journal of Chemical and Engineering Data 12, 326 (1967).

[35] J. G. Speight et al., Lange’s handbook of chemistry (McGraw-Hill New York, ADDRESS, 2005), Vol. 1.

[36] A. Klamt, F. Eckert, J. Reinisch, and K. Wichmann, Journal of Computer-Aided Molecular Design 1 (2016).

[37] W. K. Hastings, Biometrika 57, 97 (1970).

